# Transcription attenuation in synthetic promoters in tandem formation

**DOI:** 10.1101/2023.12.05.569149

**Authors:** Vatsala Chauhan, Ines S. C. Baptista, Rahul Jagadeesan, Suchintak Dash, Andre S. Ribeiro

## Abstract

Closely spaced promoters are ubiquitous in prokaryotic and eukaryotic genomes. How their structure and dynamics relate remains unclear, particularly for tandem formations. To study their transcriptional interference, we engineered two pairs and one trio of synthetic promoters in non-overlapping, tandem formation, in single-copy plasmids. From *in vivo* measurements in *E. coli* cells, we found that promoters in tandem formation have attenuated transcription rates. The attenuation strength can be widely fine-tuned by the promoters’ positioning, natural regulatory mechanisms, and other factors, including the antibiotic rifampicin, which hampers RNAP promoter escape. From this, and supported by in silico models, we concluded that the attenuation emerges from premature terminations generated by collisions between RNAPs elongating from upstream promoters and RNAPs occupying downstream promoters. Moreover, we found that these collisions can cause one or both RNAPs to fall-off. The broad spectrum of possible, externally regulated, attenuation strengths in synthetic tandem promoters should make these structures valuable internal regulators of future synthetic circuits.

## INTRODUCTION

Closely spaced promoters in convergent, divergent, and tandem geometries are widely present in living organisms, including *Escherichia coli*^1–7^. They are known to have high conservation levels^2,8^, but it remains unclear what are their selective advantages.

Provided that the elongation regions of two genes overlap, RNAPs starting from their promoters can be expected to interact. Studies reported^9^ that promoters in convergent formation have weakened RNA production, due to collisions between the RNAPs elongating from one TSS and RNAPs bound to the other. Meanwhile, in promoters in tandem formation, RNAPs bound to one promoter can block (at least transiently) the RNAPs elongating from the other promoter^10^. Moreover, natural genes controlled by tandem promoters that overall, i.e. whose transcription start sites (TSSs) are closer than ∼35 bps, have significantly weaker expression rates (on average) than more distanced ones^11^. Likely, if overlapping, when one RNAP occupies one TSS, it occludes the other TSS from RNAP binding.

It should be possible to use the interactivity between RNAPs of promoters closely spaced, in tandem formation, as a means to engineer synthetic genes with fine-tuned dynamics. Models have explored the potential dynamics of promoters in tandem formation as a function of properties such as the component promoters’ strength^11,12^. However, empirical validation is largely lacking and the potential influence of other parameters (e.g., transcription factor regulation) remains largely unexplored. Also, empirical data is lacking on the outcome of collisions between elongating RNAPs and RNAPs bound to promoters, and how it could be regulated (e.g., by tuning promoter escape rates).

Here, we studied interference between promoters in tandem formation. For this, we engineered synthetic constructs of promoters in tandem formation using, as building blocks, three genetically modified natural promoters: P_LacO3O1_, P_tetA_, and P_BAD_^13–15^. The internal composition of our synthetic constructs is illustrated in Figure 1. In all constructs, the promoter(s) control the expression of an mCherry protein to track their transcription dynamics. We used the constructs to study interference as a function of the regulation state of the component promoters, and when subject to an antibiotic that directly interferes with transcription initiation^16^.

**Figure 1:**
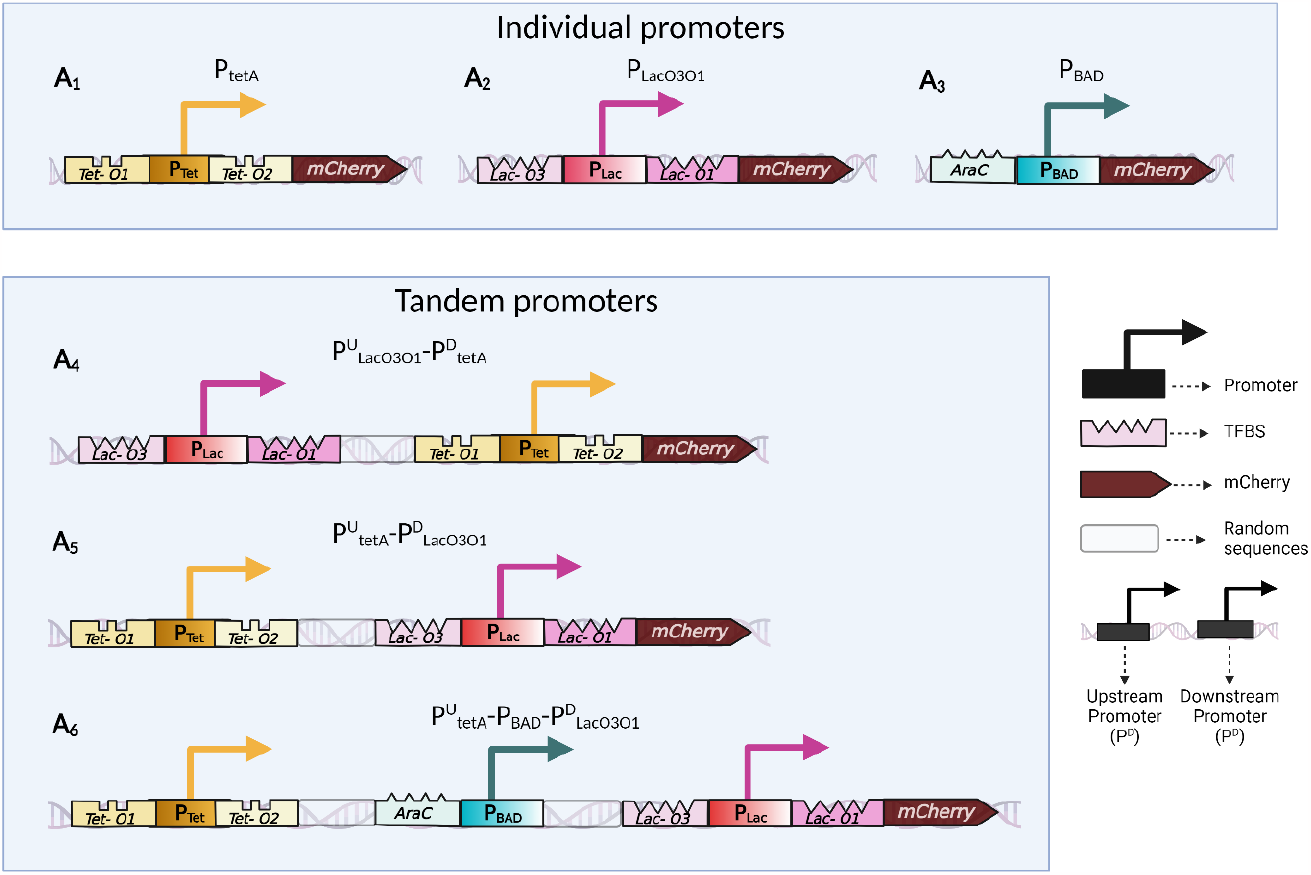
Schematic representation of the synthetic promoters in tandem formation along with their transcription factor binding sites. Bioparts A_1_-A_6_ were each inserted into single-copy pBAC plasmids, respectively. Each biopart is followed by an mCherry coding region. Bioparts A_1_, A_2_, and A_3_ are the individual promoter constructs. Bioparts A_4_, A_5_, and A_6_ are the constructs with (pairs or trios of) tandem promoters. The component promoters in individual formation are P_tetA_, P_LacO3O1_, and P_BAD_, respectively. The two dual synthetic promoters in tandem formation are P^U^_LacO3O1_-P^D^_tetA_ and P^U^_tetA_-P^D^_LacO3O1_ respectively, (U and D stand for upstream and downstream respectively). Finally, the trio of promoters in tandem formation is P^U^_tetA_-P_BAD_-P^D^_LacO3O1_.

From Figure 1, the fluorescence probes of the tandem constructs allow tracking the overall dynamics of the component promoters (Figures 1A_4_ to 1A_6_), but not the dynamics of each component promoter independently. For that, we used individual promoter constructs (Figures 1A_1_ to 1A_3_). From here onwards, we referred to the constructs carrying promoters in tandem formation as “tandem promoters” (e.g., tandem promoters “P^U^_tetA_-P^D^_LacO3O1_”). Meanwhile, we referred to the constructs carrying a single promoter as “individual promoters” (e.g., “individual promoter P_tetA_”).

## RESULTS

The promoters used to build the tandem promoters have been profusely studied. P_LacO3O1_ was engineered from the natural Lac promoter of the lactose operon in *E. coli*^17^, by removing the operator O_2_^18,19^. This decreases by 2-to 3-fold the repression strength of the wild-type tetrameric Lac repressor (LacI)^18^. The homotetrameric Lac repressor protein represses by binding the DNA operator sequences^20^. The binding forces DNA loops^21,22^ that make the promoter less accessible^23^. Contrarily, IPTG, a structural analog of the natural inducer Lactose^24^, indirectly induces P_LacO3O1_ by binding to LacI, which hampers LacI’s ability to bind to the promoter^20^.

The second promoter used, P_tetA_, was extracted in its natural from the Tet operon in *E. coli*, where it controls the expression of the tetA gene^25^. The Tet operon is involved in tetracycline resistance and can self-repress due to carrying a second gene, tetR, coding for TetR^26^. This protein binds to the operator sites of the promoters of tetA and tetR and prevents their transcription^27^. Meanwhile, tetracycline binds to TetR hampering its ability to bind to the DNA, thus enabling the expression of tetA and tetR^28^. aTc, an analog of tetracycline, can induce P_tetA_ by the same process^29^.

The final promoter used, P_BAD_, was also extracted in its natural form. Originally, it is a component of the L-arabinose operon of *E. coli*^30^. It can be repressed by dimers of AraC, which can form a DNA loop that blocks transcription^31^. Arabinose can induce P_BAD_ by binding to the dimeric AraC. This binding changes the conformation of AraC, which breaks the DNA loop, allowing RNAP to bind to P_BAD_^32^.

A recent work used similar promoters to study the dynamics of genes in convergent, divergent, and tandem formations^33^. However, unlike in our constructs, the elongation regions were separated by Rho independent hairpin loops. Thus, the RNAPs transcribing from one promoter should not collide with RNAPs transcribing from the other promoter. Among other, they used the constructs to study the influence of supercoiling buildup on closely spaced genes.

Next, we describe the assembly of the genetic constructs. Afterwards, we study their dynamics and show that, in tandem formation, their strength is reduced due to collisions between RNAPs leading to premature transcription terminations. We also show that this phenomenon can be fine-tuned by inducers of the regulatory mechanisms of the component promoters and by a transcription-targeting antibiotic. Moreover, we use a stochastic model to show that the phenomena identified suffices to explain the observed dynamics of the promoters in tandem formation. In the end, we discuss how our constructs may become valuable components of future synthetic genetic circuits.

### Assembly of the synthetic, non-overlapping tandem promoters in single-copy plasmids

We designed the tandem constructs using Snapgene (GSL Biotech) and assembled them at Integrated DNA Technology, Iowa, U.S.A. Next, we introduced them into single-copy plasmids (pBAC)^34^. The original sequences of P_LacO3O1,_ P_tetA,_ and P_BAD_ (Methods Section “Bacterial strains, growth conditions, induction, and antibiotic”) were not altered. We distanced the transcription start sites (TSSs) of the pairs of promoters in tandem formation by 150 bps (Figures 1A_4_ and 1A_5_), while the trio of promoters were separated by 200 bps, each (Figure 1A_6_). For that, we introduced random sequences between the promoters generated using Random.org. Snapgene confirmed the absence of sequences known to lead to the formation of secondary RNA structures (e.g., hairpin loops) as well as of sequences coding for translational products.

In addition to the tandem promoters (“biopart”), each plasmid has a sequence coding for a fluorescent protein, mCherry^35^ (plasmid accession number KX264176.1^34,35^)), immediately downstream of the biopart. The DNA code includes a strong RBS, to ensure strong translation rates to robustly track promoter activity. Moreover, the maturation time of mCherry (FPbase ID: ZERB6) is 15 minutes^36^, to closely report RNA production rates. We also produced three control strains, each carrying one individual promoter (P_LacO3O1,_ P_tetA,_ and P_BAD_, respectively) (Figures 1A_1_-A_3_).

Since all constructs have the same RNA degradation rate and mCherry translation and degradation rates, differences in fluorescence intensities between strains should be caused mostly by differences in overall transcription rates alone. To best ensure this, small differences in growth rates between strains were accounted for (next section). Finally, since some tetracycline derivatives exhibit fluorescence^37^, we tested if the inducer aTc could interfere with the measurements. However, within the range of concentrations used, aTc did not significantly affect cell fluorescence (Figure S1).

### Cell morphology and physiology. Variability in single cell expression levels

We searched for potential physiological and morphological differences between strains that could affect the constructs’ expression. First, we monitored cell growth rates of all strains in all measurement conditions studied throughout the manuscript (Figure 2A). Strains carrying the plasmids have a slightly slower growth rate than the wilt type, WT, strain (8% weaker). For comparison, rifampicin (5 μg/mL) reduced growth by much more (41%). Given the existence of differences, from here onwards, we accounted for them by normalizing mean population expression levels obtained by spectrophotometry using data on cell populations sizes using data from O.D._600_ measurements (Methods Section “Spectrophotometry”).

**Figure 2:**
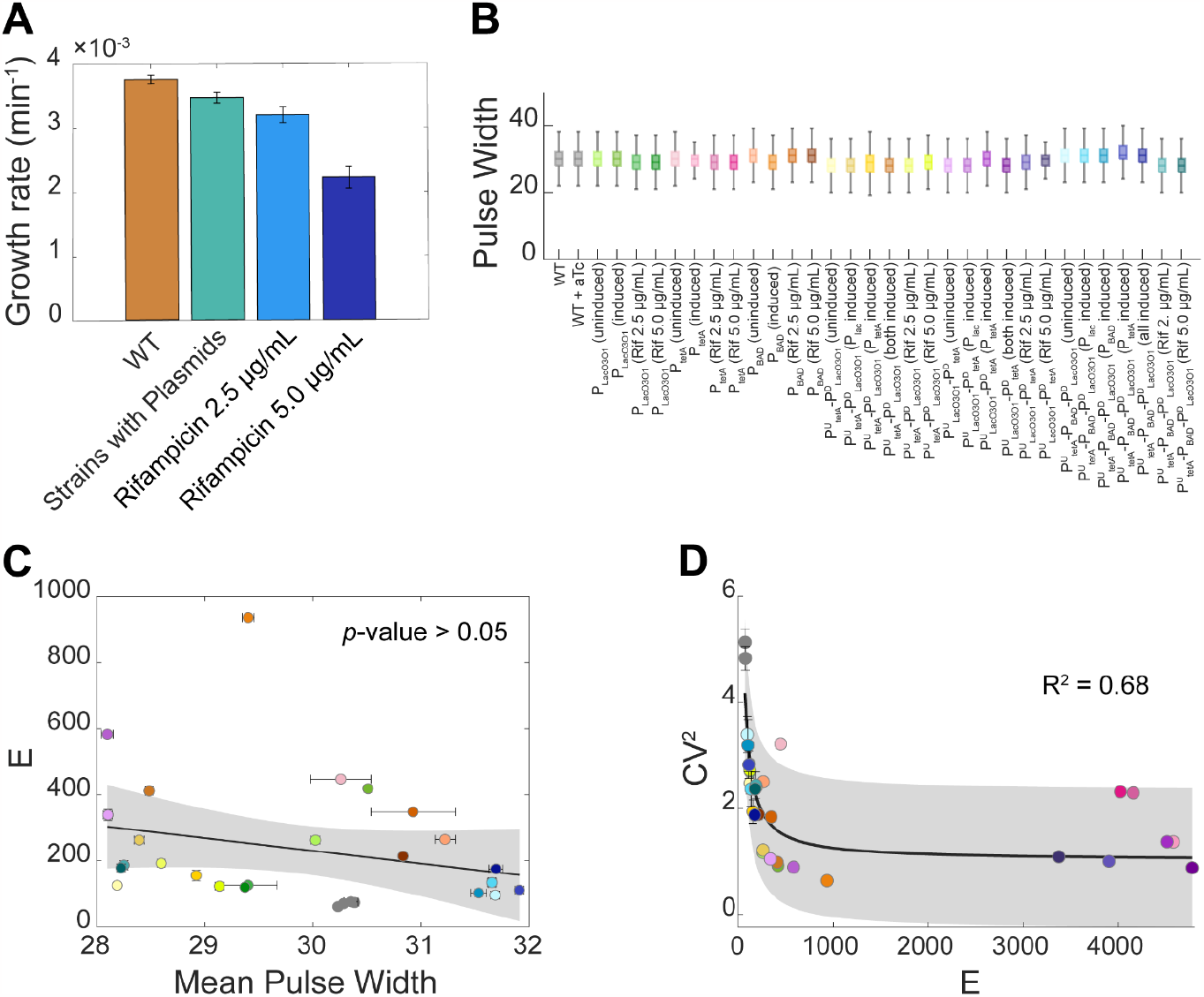
**(A)** Average growth rates (O.D._600_ measurements by spectrophotometry). **(B)** Boxplots of the distributions of single-cell pulse width, used as a proxy for cell size. The outliers (values higher or lower than 1.5·IQR, where IQR is the interquartile range) are not shown. **(C)** Scatter plot of the mean single-cell fluorescence, *E*, and the mean pulse width. Also shown are the best-fitting line and corresponding *p*-value, after excluding outliers (not shown) (Methods section” Fitting and statistical analysis”). The correlation is not significant at the 5% significance level, even when including outliers (Figure S2). **(D)** Scatter plot between the CV^2^ and the mean single-cell fluorescence intensities (*E*). Also shown is the best-fitting curve and the corresponding R^2^ (Methods section” Fitting and statistical analysis”). The shaded area represents the 95% confidence interval. The equation and coefficient values of the fitted curve are shown in Supplementary Table S4. Finally, the error bars in **(A), (C), (D)** correspond to the standard error of the mean of 3 biological replicates.

Next, we observed cell sizes using pulse width from flow cytometry as a proxy^38^. The pulse width differs little between strains and conditions (Figure 2B). In agreement, we could not find a significant correlation between mean single-cell protein expression levels (*E*) and mean single-cell pulse widths (Figure 2C). Thus, gene expression intensities obtained by flow cytometry were not normalized by cell sizes.

Finally, we searched for correlations (in flow cytometry data) between mean single-cell expression levels and cell-to-cell variability (as measured by the squared coefficient of variation CV^2^). The high R^2^ value of a curve (equation 1) accounting for a noise floor “*n*”^39^ (Figure 2D) suggests that the constructs follow the natural genome-wide behavior^40,41^.

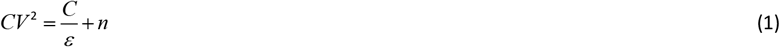

### Promoters in tandem formation have attenuated overall expression rates

Next, we studied the dynamics of the dual synthetic tandem promoters P^U^_LacO3O1_-P^D^_tetA_ and P^U^_tetA_-P^D^_LacO3O1_ from single-cell fluorescence data under three induction schemes: (i) upstream promoter induced; (ii) downstream promoter induced; and (iii) both promoters induced. For comparison, we also studied the individual promoters P_LacO3O1_ and P_tetA_, when and not induced.

The mean, standard deviation, CV^2^, skewness and kurtosis of the distributions of single-cell expression levels are shown in Supplementary File 1.

From the data, first, we quantified the attenuation in the expression of the promoters, when in tandem formation, when compared with the component promoters. Let *T* stand for the tandem promoters, *U* stand for the individual upstream promoter, and *D* stand for the individual downstream promoter. We define “absolute attenuation”, *α*_*A*_, in the tandem promoters as:

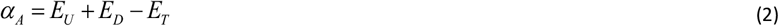

From Figure 3, in both tandem constructs under all induction schemes, *α*_*A*_ > 0. Thus, in general, placing these promoters in tandem formation reduces overall protein expression rates.

**Figure 3:**
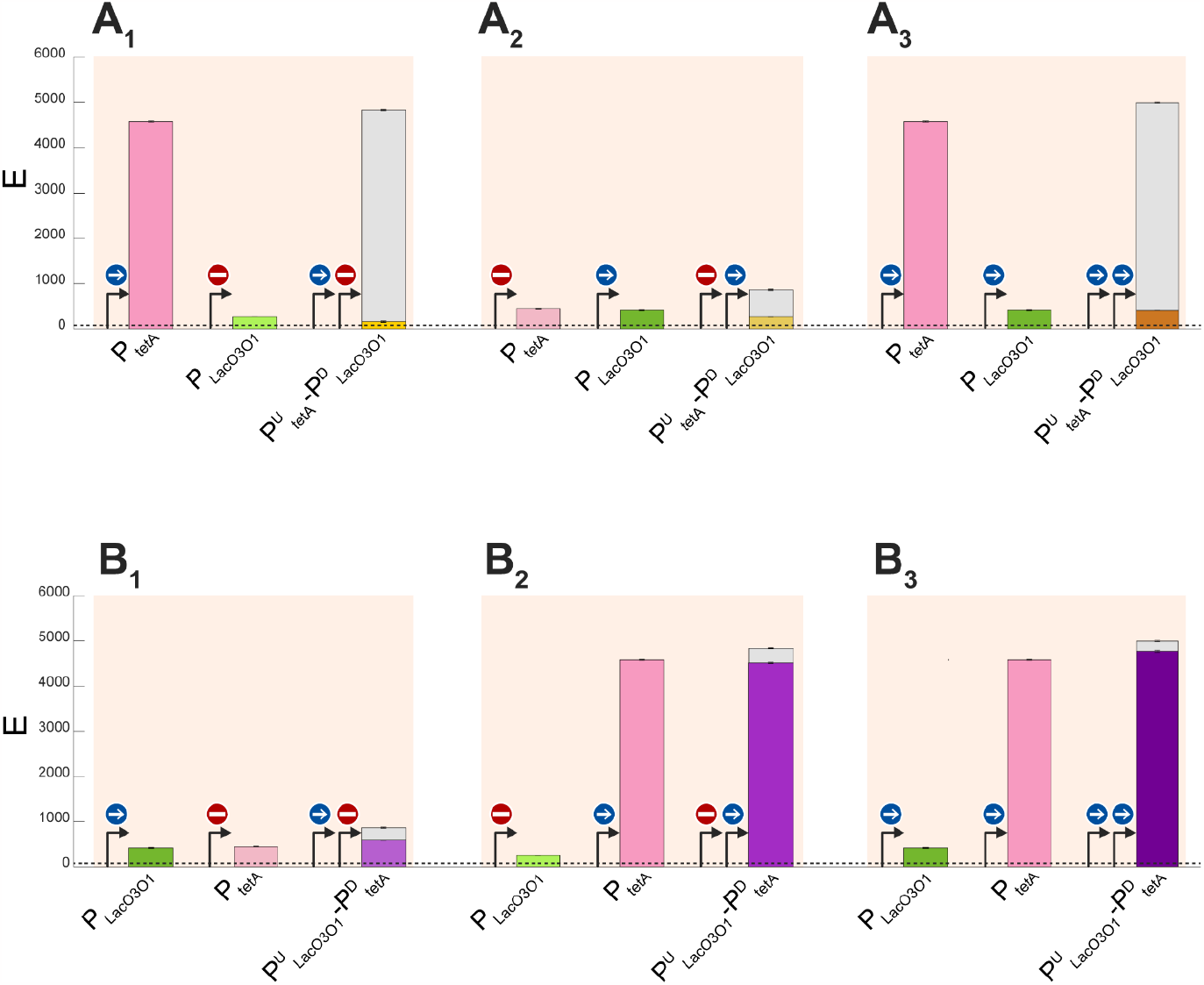
Mean expression intensity (*E*) of the tandem and the individual promoters, under various induction schemes. The blue sign with a straight white arrow stands for “fully induced”, while the red “Do Not Enter” sign stands for “fully repressed” **(A**_**1**_**-A**_**3**_) P_tetA_, P_LacO3O1,_ and P^U^_tetA_-P^D^_LacO3O1_, respectively. **(B**_**1**_**-B**_**3**_) P_tetA_, P_LacO3O1_, and P^D^_LacO3O1_-P^U^_tetA_, respectively. In all plots, the height of the gray bars equals the absolute attenuation, *α*_*A*_, defined in equation (2). I.e., they correspond to the “loss” in mean expression intensity of two promoters, when placing them in tandem formation. The error bars correspond to the standard error of the mean of 3 biological replicates. The dashed horizontal line near 0 marks the autofluorescence intensity of WT cells. The expression levels of the individual promoters, P_tetA_ and P_LacO3O1_, are shown more than once to facilitate comparing the tandem constructs with the component promoters in individual formation.

Meanwhile, for P^U^_tetA_-P^D^_LacO3O1_ alone: *E*_*T*_ < *E*_*U*_ and *E*_*T*_ < *E*_*D*_ (Figures 3A_1_-3A_3_). I.e., placing the weaker promoter downstream heavily reduced the RNA production rate of the upstream promoter, causing P^U^_tetA_-P^D^_LacO3O1_ expression to be weaker than the downstream promoter alone.

Conversely, for P^U^_LacO3O1_-P^D^_tetA_, in one condition we find that: *E*_*T*_ > *E*_*U*_ and *E*_*T*_ < *E*_*D*_ (Figure 3B_2_). Interestingly, we also observe that when P_LacO3O1_ is induced: *E*_*T*_ > *E*_*U*_ and *E*_*T*_ > *E*_*D*_ (Figures 3B_1_ and 3B_3_). I.e., the tandem promoters P^U^_LacO3O1_-P^D^_tetA_ express more strongly than the individual downstream promoter.

From the above, we interpret the positive absolute attenuation in both constructs in all conditions (Figures 3A_1_-3A_3_ and 3B_1_-3B_3_) as evidence that many RNAPs elongating from the upstream promoter are prematurely terminated (fall-off). When the downstream promoter is repressed, the fall-offs are likely due to the repressors’ binding (Figure 4C). Conversely, when the downstream promoter is active, the fall-offs are likely due to collisions between the elongating RNAP from the upstream promoter with an RNAP bound to the downstream promoter (Figure 4D).

**Figure 4:**
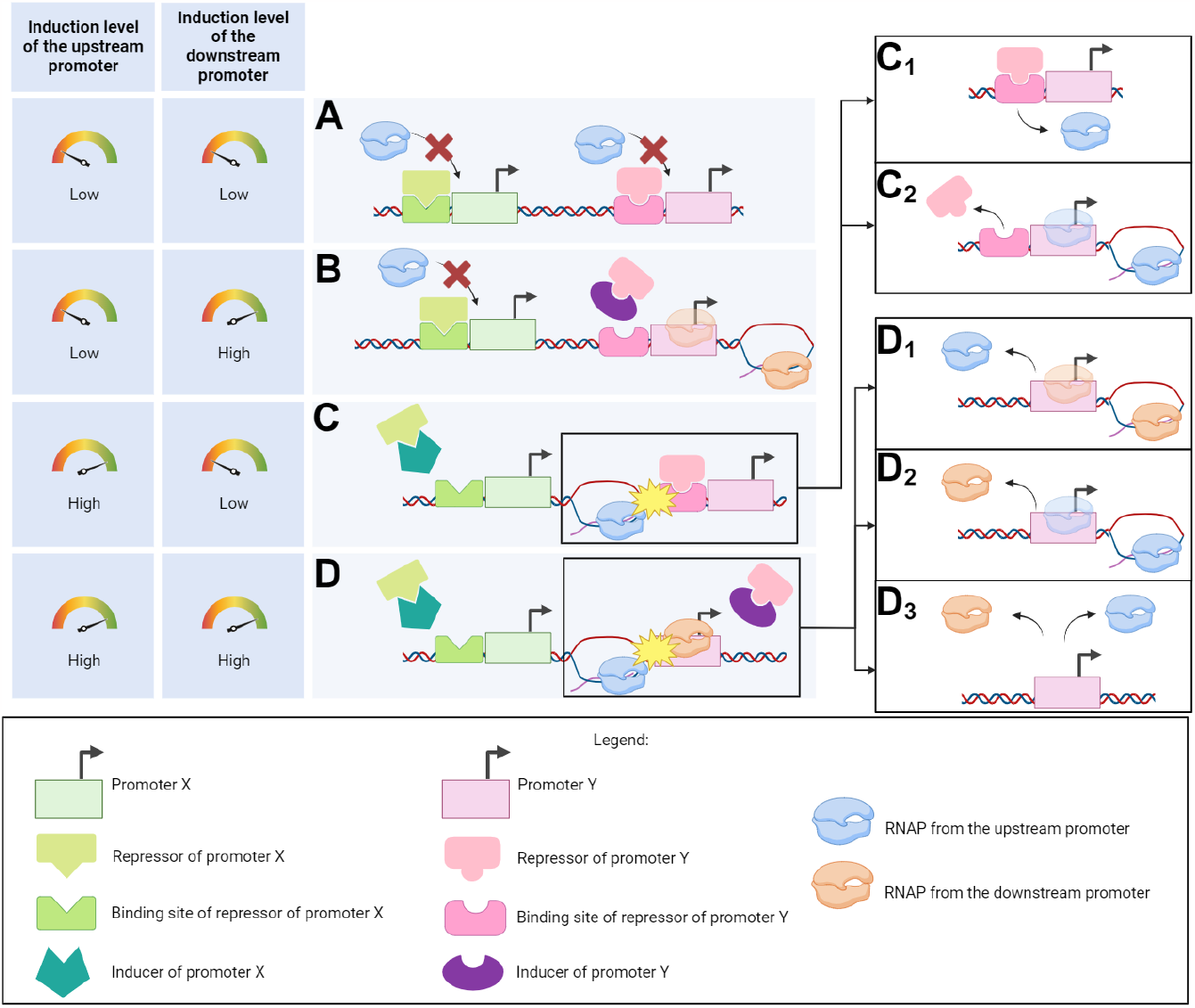
Illustration of transcription interference events generating attenuation in tandem promoters under different induction schemes. **(A)** Weak-to-no induction of both the upstream and the downstream promoters. **(B)** Weak-to-no induction of the upstream promoter along with high induction pf the downstream promoter. **(C)** High induction of the upstream promoter and weak-to-no induction of the downstream promoter. **(D)** High induction of both the upstream and downstream promoters. In **(C)** collisions between RNAPs elongating from the upstream promoter and the repressor at the downstream promoter are likely to occur. Those collisions can cause **(C**_**1**_) fall-off of the elongating RNAP, or **(C**_**2**_) fall-off of the repressor bound to the operator site. Similarly, in **(D)** the RNAP elongating from the upstream promoter can collide with the RNAP occupying the downstream promoter. This causes **(D**_**1**_) the elongating RNAP to fall-off, **(D**_**2**_) the RNAP at the downstream promoter to fall-off, or **(D**_**3**_) both RNAPs to fall-off.

Moreover, we interpret the existence of a tandem construct with weaker activity than either of its component promoters (Figures 3A_1_-3A_3_) as evidence that several collisions cause fall-offs of both the RNAP bound to the downstream promoter as well as the elongating RNAP (Figure 4D_3_). If this never occurred, the tandem promoters should be at least as strong as the downstream promoter alone.

Finally, we interpret the existence of a tandem construct with stronger activity than the individual downstream promoter (Figures 3B_1_ and 3B_3_) as evidence that, in some tandem promoters, neither the repressors (Figure 3B_1_), nor RNAPs initiating transcription (Figure 3B_3_) at the downstream promoter, can block all RNAPs elongating from the upstream promoter (Figures 4C_2_ and 4D_2_).

### Attenuation in synthetic tandem constructs can be fine-tuned by induction of the component promoters

We investigated if the attenuation can be fine-tuned, i.e., is sensitive to external regulation of the activity of the component promoters. We tested in P^U^_tetA_-P^D^_LacO3O_, since it exhibited stronger attenuation. To compare attenuations as a function of inductions strength, we defined relative attenuation, *α*_*R*_, as:

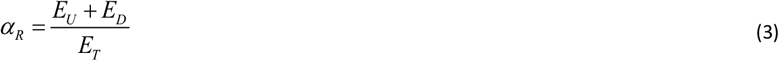

We started with a fully induced P^D^_LacO3O1_ and then gradually induced P^U^_tetA_. Visibly, P^U^_tetA_-P^D^_LacO3O1_ is less induced than P_tetA_ alone (Figure 5A_1_), and *α*_*R*_ increases with induction of the upstream promoter (Figure 5A_2_). This can be explained by an increase in the fraction of RNAPs from the upstream promoter that fail to complete elongation. Such fall-offs could be explained by the transient occupation of the (active) downstream promoter by RNAPs initiating transcription.

**Figure 5:**
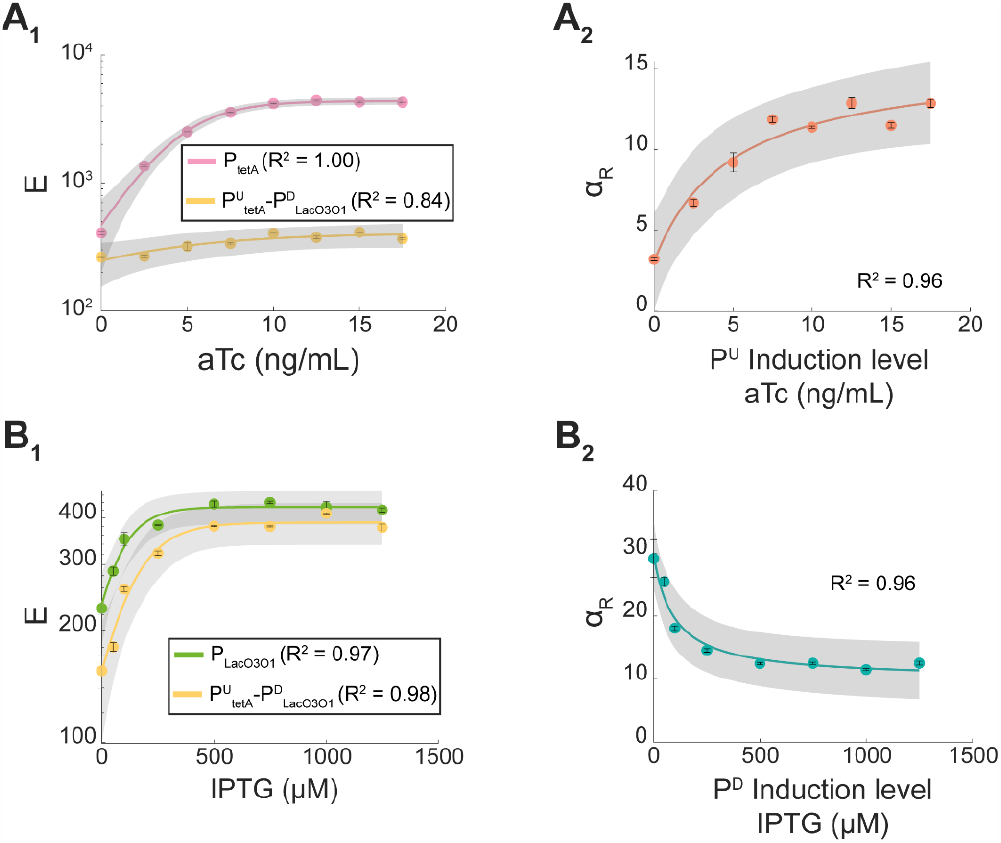
**(A**_**1**_) Mean single-cell expression level (*E*) of P_tetA_ and of P^U^_tetA_-P^D^_LacO3O1_, when inducing P_tetA_. In all conditions, P^D^_LacO3O1_ is fully induced. **(A**_**2**_) Attenuation of the tandem promoters with increasing induction of the upstream promoter. **(B**_**1**_) Mean single-cell expression level (*E*) of P_LacO3O1_ and of P^U^_tetA_-P^D^_LacO3O1_, when inducing P_LacO3O1_. In all conditions, P^U^_tetA_ is fully induced. **(B**_**2**_) Attenuation of the tandem promoters with increasing induction of the downstream promoter. In all plots, the error bars correspond to the standard error mean of 3 biological replicates. Also shown are best fitting curves and the corresponding R^2^ (Methods section” Fitting and statistical analysis”). The shaded areas represent the 95% confidence intervals. The equation and coefficient values of the fitted curve are shown in Supplementary Table S4. Note that the vertical axes of (B_1_) and (A_1_) are in logarithmic scale.

Next, we started with a fully induced P^U^_tetA_ and gradually induced the downstream promoter, P^D^_LacO3O1_. Again, the individual promoter (P_tetA_) is more strongly induced than the tandem construct, implying the existence of attenuation (Figure 5B_1_). Nevertheless, contrary to before, *α*_*R*_ decreased with induction (Figure 5B_2_). This is expected since repressors bound to the DNA are expected to be more efficient in blocking RNAPs elongating from the upstream promoter than RNAPs occupying the downstream promoter. Consequently, as the repression mechanism of the downstream promoter is inactivated, more RNAPs from the upstream promoter can complete elongation.

Given all of the above, we conclude that *α*_*R*_ can be fine-tuned by the regulatory mechanisms of the component promoters in diverse ways.

#### Adding another promoter in tandem formation enhances attenuation

We next introduced an additional promoter, P_BAD_, in between two tandem promoters (P^U^_tetA_-P^D^_LacO3O1_). From this resulted the trio of tandem promoters: P^U^_tetA-_P_BAD_-P^D^_LacO3O1_. To study its attenuation, we defined, where *M* stands for “Middle”:

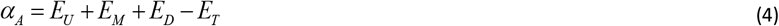

Expectedly, P^U^_tetA-_P_BAD_-P^D^_LacO3O1_ activity is also weaker than the sum of individual promoter activities. I.e., in all induction schemes (Figures 6A_1_-A_4_), *α*_*A*_ > 0.

**Figure 6:**
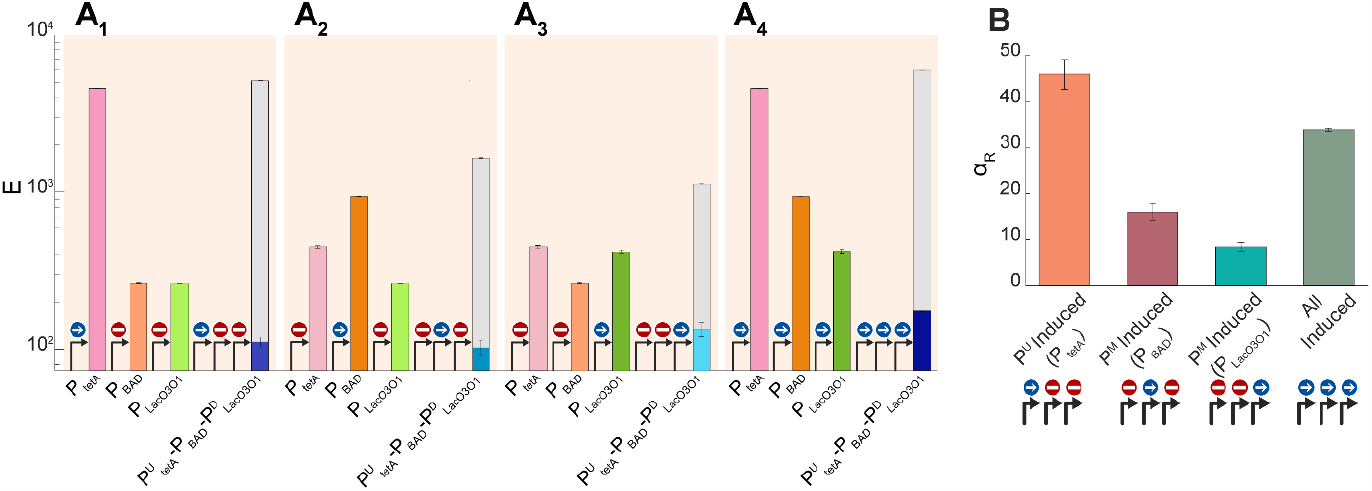
**(A_1_-A_4_)** Mean expression intensities (*E*, in logarithmic scale) of the trio tandem promoters (P^U^_tetA-_P_BAD_-P^D^_LacO1O3_) and of each individual promoter. We tested inducing only the most upstream promoter (A_1_), inducing only the middle promoter (A_2_), inducing only the most downstream promoter (A_3_), and inducing all promoters at the same time (A_4_). The gray bars measure the absolute attenuation, *α*_*A*_, defined in equation (4). **(B)** Relative attenuation, *α*_*R*_ (Equation 5), of the trio of tandem promoters under various induction schemes. In all plots, the error bars correspond to the standard error of the mean of 3 biological replicates.

Conversely, unlike the dual tandem promoters, the *E*_*T*_ of the trio of tandem promoters is smaller than *E*_*U*_, *E*_*M*_, and *E*_*D*_, alone, in all induction schemes (Figure 6A): *E*_*T*_ < *E*_*U*_, *E*_*M*_, *E*_*D*_. This is evidence for the occurrence of many collisions between RNAPs elongating from the upstream promoters with repressors (causing the fall-off of the RNAP) and with RNAPs occupying the downstream promoters (causing the fall-offs of both colliding RNAPs).

The strongest *α*_*R*_ (Equation 5) still occurs when the most upstream promoter is the only one induced (Figure 6B), suggesting that, as expected, collisions between elongating RNAPs and repressors bound to the DNA usually cause the collision of the RNAP. Interestingly, *α*_*R*_ is also high when all promoters are induced. The simplest explanation is that numerous RNAPs elongating from the upstream promoters are prematurely terminated by collisions with RNAPs occupying the downstream promoters, and that several collisions cause both colliding RNAPs to fall-off.

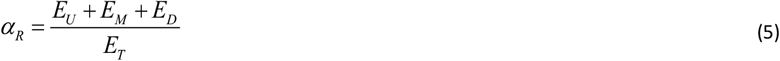

### Attenuation is enhanced when hampering promoter escape by adding rifampicin

Above, we argued that several RNAPs are prematurely terminated by collisions of RNAPs elongating from upstream promoters with RNAPs sitting at downstream promoters. If this holds true, increasing the fraction of time spent by RNAPs occupying a downstream promoter should increase α_R_.

To test this, we subjected cells to rifampicin (Methods section “Fitting and statistical analysis”). This antibiotic binds to the β sub-unit of RNAP^42^. When bound, the RNAP cannot escape beyond 2-3 nucleotides away from the TSS^43,44^. Thus, rifampicin should not only reduce the activity of each component promoter, but also increase *α*_*R*_, due to increasing the occupancy time of RNAPs at the downstream promoter (illustrated in Figure 7A). Other direct effects on gene expression are not expected since, e.g., rifampicin does not affect stable transcription elongation complexes^45,46^.

**Figure 7:**
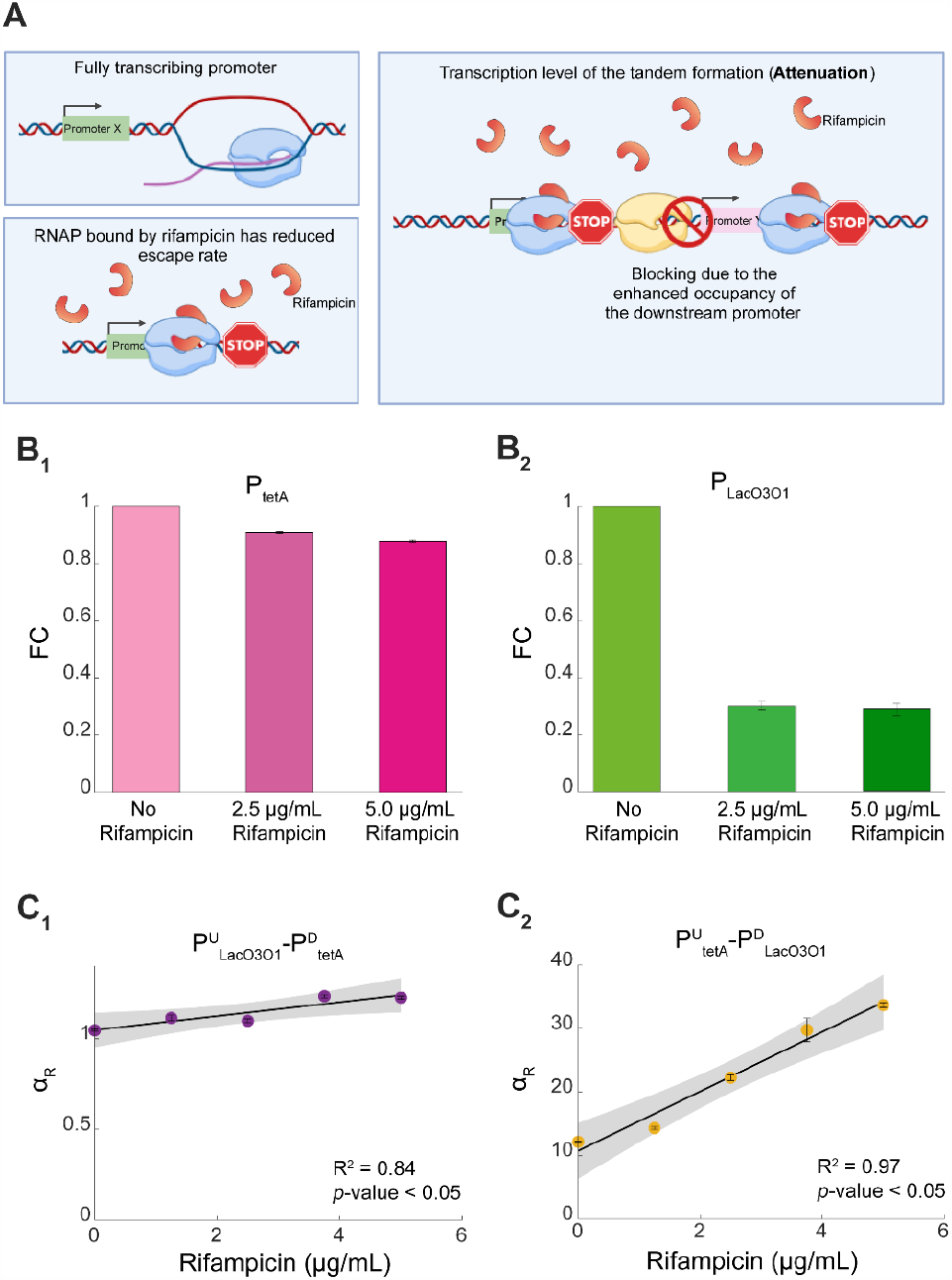
Effects of rifampicin on the attenuation in tandem promoters. **(A)** Illustration of how rifampicin reduces the activity of an individual promoter, as well as of promoters in tandem formation. The latter should be further affected by enhanced attenuation. **(B**_**1**_**-B**_**2**_) Fold-change in mean expression levels (*FC*) of P_tetA_ and P_LacO3O1_ due to rifampicin. **(C**_**1**_**-C**_**2**_) Relative attenuation (*α*_*R*_) of P^U^_tetA_-P^D^_LacO3O1_ and P^U^_LacO3O1_-P^D^_tetA_, when subject to rifampicin. The error bars correspond to the standard error of the mean of 3 biological replicates. Shown are the best-fitting lines and their *p*-value and R^2^ (Methods section “Fitting and Statistical Analysis”). The shaded areas represent the 95% confidence interval. The equation and coefficient values of the fitted lines are shown in Supplementary Table S4.

As expected, rifampicin reduced the activity of both P_tetA_ as well as P_LacO3O1_ in individual formations (Figures 7B_1_ and 7B_2_, respectively). P_LacO3O1_ was (relatively) more affected than P_tetA_, for unknown reasons (also visible in spectrophotometry data in Figure S3). Also, for unknown reasons, P_tetA_ activity reduction was approximately linear with rifampicin concentration, while P_LacO3O1_ had a sharp initial decrease in expression, but further increasing rifampicin concentration did have significant additional effects.

Most importantly, we also observed that rifampicin gradually increased *α*_*R*_ in both tandem constructs (Figures 7C_1_ and 7C_2_), as hypothesized. Noteworthy, the higher sensitivity of P_LacO3O1_ to rifampicin explains why *α*_*R*_ of P^U^_tetA_-P^D^_LacO3O1_ changed the most (Figures 7C_1_ and 7C_2_).

These results exemplify how the attenuation in tandem promoters can be controlled by external regulation and, with that, produce different (yet predictable) gene expression dynamics.

### Model of transcription attenuation of promoters in tandem formation due to RNAP collisions and consequent fall-offs can explain the empirical data

Above, we observed that the tandem promoters always exhibit attenuation. In P^U^_tetA_-P^D^_LacO3O1_, the attenuation was strong, in general, making expression levels lower than in the individual downstream promoter. Instead, in P^U^_LacO3O1_-P^D^_tetA_, the attenuation was weak, allowing, in some conditions, almost as strong expression as the sum of expressions of the individual component promoters. In both tandem promoters, attenuation was present when the downstream promoter was activated, as well as when it was repressed, in which case it was stronger.

In this section, we propose a general stochastic model for tandem promoters. Shortly, the model includes a two-step transcription initiation process at each promoter, to account for promoter occupancy by RNAPs. Repressors can also occlude promoters, while inducers inactivate repressors. Also, RNAs degrade. Finally, we model collisions between RNAPs, as well as between RNAPs and repressors occluding promoters. Critically, we assume that the collisions between RNAPs can cause fall-offs of one or both RNAPs. Meanwhile, collisions between RNAPs and repressors always cause the RNAPs to fall-off. A complete model description is provided in Supplementary Section S1. Using this model, we show that various degrees of attenuation observed above can be explained by fall-offs emerging from RNAP-RNAP and RNAP-repressor collisions.

First, we tested whether increasing the time that the downstream promoter is occupied by initiating RNAPs increases attenuation (due to increased rate of collisions and subsequent fall-offs). For this, we decreased the rate of RNAP escape from the downstream promoter (*k*_*esc*_^*D*^) (similar to the effects of increasing rifampicin, Figures 7C_1_-C_2_). This increased the mean amount of time that the downstream promoter is occupied by RNAPs (Figure 8A). This, in turn, increased linearly the relative attenuation (Figure 8B), as in Figures 7C_1_-C_2_. Also, in agreement with the empirical data (Supplementary file 1), it decreased E_D_ (Figure 8C) and E_T_ (Figure 8D).

**Figure 8:**
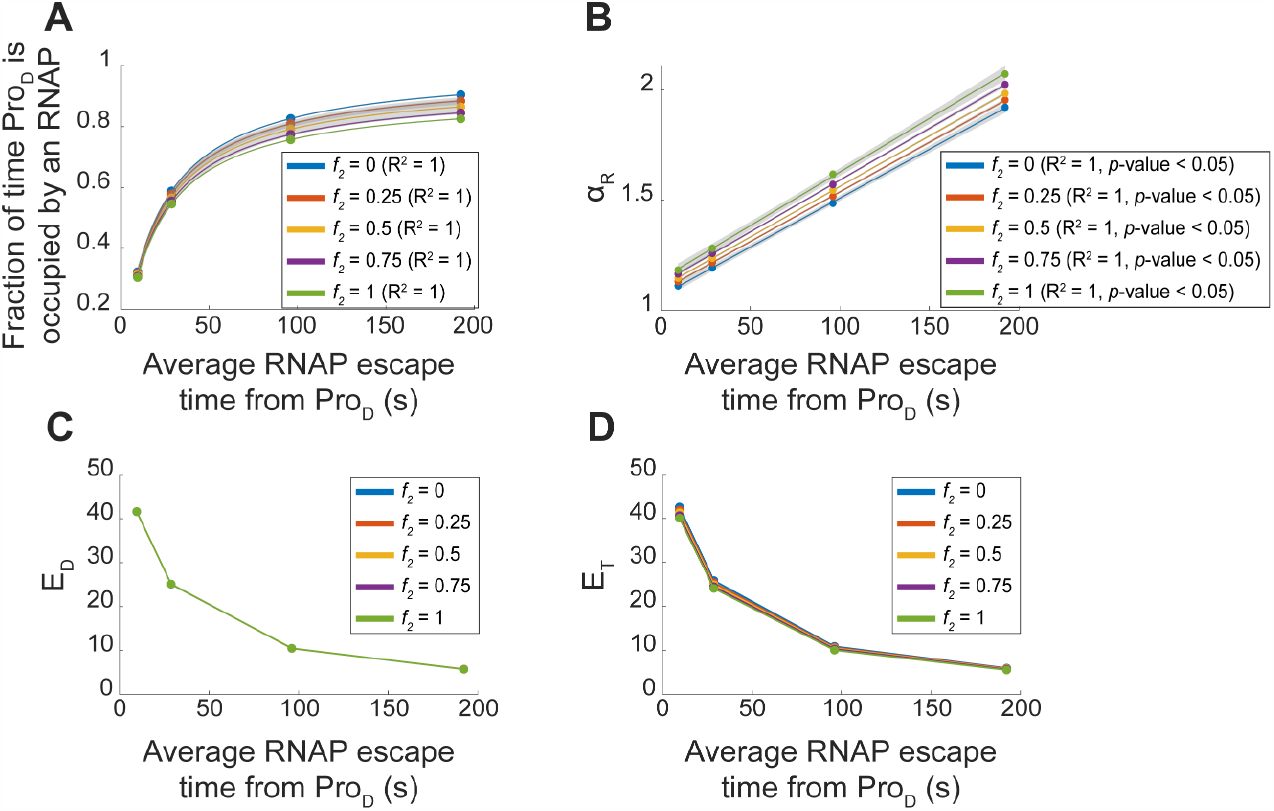
Estimations using the stochastic models of the effects of increasing the average time for RNAPs to escape the downstream promoter (Pro_D_), by decreasing *k*_*esc*_^*D*^, as well as of changing the frequency with which RNAP collisions cause both RNAPs to fall-off, instead of only one falling-off. **(A)** Fraction of time that the downstream promoter is occupied by an RNAP as a function of the inverse of the RNAP escape rate. Note that the latter is affected also by collisions between RNAPs. **(B)** Relative attenuation (*α*_*R*_) in tandem promoters as a function of the average RNAP escape times from downstream promoters. **(C-D)** Expression levels of the individual downstream promoter (E_D_) and of the tandem promoters (E_T_) as a function of the average RNAP escape times from the downstream promoter. In all plots, each colored line represents a different relative frequency (*f*_*2*_) of both RNAPs falling-off upon colliding (as opposed to only one RNAP falling-off). (A) and (B) also show the best fitting functions along with their R^2^ values (and *p*-values in case of linear fits) (Methods section” Fitting and statistical analysis”). The shaded areas represent the 95% confidence intervals. The equation and coefficient values of the fitted lines are shown in Supplementary Table S4.

Next, we tested increasing the frequency *f*_*2*_ with which RNAP-RNAP collisions cause both RNAPs to fall-off (instead of only one RNAP falling-off). We observed, first, a small decrease in the fraction of time that the downstream promoter is occupied by RNAPs (Figure 8A), as expected from the increased fall-offs of the initiating RNAPs. In agreement, we observe increased *α*_*R*_ with increased *f*_*2*_ (Figure 8B). Meanwhile, the quantitative relationships between E_D_ and E_T_ with the average RNAP escape rates (Figure 8C-8D) are not influenced (since both change accordingly).

We also observed that, for higher frequencies of double RNAP fall-offs, E_T_ can become slightly weaker than E_D_ (Supplementary Figure S4A). We explored this by tuning the binding and escape rates of RNAPs to Pro_U_ and Pro_D_. From Supplementary Figure S4B, there is a wide range of parameter values (Supplementary Table S3) for which E_T_ can be considerably weaker than E_D_ (in agreement with the empirical data in Figures 3A_1_-3A_3_).

Finally, we tested with the model if the induction-repression mechanisms can increase and decrease *α*_*R*_ with increasing induction, depending on whether it is the upstream or the downstream promoter that are initially repressed. Supplementary Figure S5 supports that this is possible. Overall, we conclude that the model can explain all behaviors observed in the two tandem constructs.

## DISCUSSION

Above, we found that premature transcription terminations of RNAPs elongating from upstream promoters and of RNAPs occupying downstream promoters can cause significant attenuation of tandem constructs. The attenuation can differ widely with the active transcription initiation dynamics of the component promoters, allowing for the expression of the tandem constructs to range from weaker than either component promoter up to similar (but never higher) than the sum of the expression dynamics of the two promoters. Moreover, the attenuation can be further externally tuned using the natural regulatory mechanisms of each promoter and/or using antibiotics targeting, e.g., transcription. This tunability should allow tandem formations to execute fine-tuned expression dynamics.

Tandem constructs dynamics were largely predictable by a simple model that only requires knowledge of the dynamics of the component promoters in individual formation, along with the relative rate of collisions that lead to two (instead of one) RNAP fall-offs. This rate should differ with the binding affinity of RNAPs or repressors to the downstream promoter and can be empirically estimated.

This model-based predictability is of significance since it remains an important and difficult challenge in synthetic biology. For example, we are yet to efficiently predict how random DNA sequences express, even in bacteria, albeit significant recent successes^47–54^.

On the contrary, knowledge of the dynamics of individual, natural promoters (e.g. in *E. coli*) is relatively easy to acquire and the data is rapidly increasing^41,55–58^. Using these increasing libraries, along with the model proposed, it should be feasible to engineer novel tandem promoters with desired, diverse dynamics. These novel constructs could then be used, e.g., as building blocks for future, more complex circuits.

As an example, genetic switches are circuits composed of two genes repressing one another. Because of this, they are expected to have two possible states: either one gene is “ON” and the other one is “OFF”, or vice-versa. One of the most promising applications of these circuits is as components of information processing circuits (e.g., for storing information). The main difficulty in achieving this is in tuning a switch to be both sufficiently sensitive to changes supposed to control their state, while also being robust in maintaining the state, if no relevant changes occur in the cell. This requires specific expression levels of both genes of the switch ^59^, and depend on promoter occupancy times by RNAP and repressors, repressor-activator binding and unbinding rates, and other parameters^60^. Finding natural promoters with the desired features or changing their sequence to achieve them is presently complex. Meanwhile, our results suggest that starting with a large library (such as^41,57^), and then use the model to find combinations of two promoters in specific states that achieve a desired dynamics is a more promising approach.

As a side note, while not observed here, we do not exclude the possibility that a few promoters in tandem formation can be stronger than the sum of the two component promoters independently. For example, this could potentially be possible if elongating RNAPs from the upstream promoter would dislodge repressors at the downstream promoter, while not falling-off themselves. Weak repressor-DNA binding affinities could make this possible. However, we do not expect it to be common in natural genomes. Nevertheless, removing repressor binding sites (or altering their sequences) could facilitate it in synthetic circuits.

In the future, we plan to explore broadly how to tune the attenuation in synthetic as well as in natural promoters in tandem formation. In addition to repression mechanisms and antibiotics hampering RNAP promoter escape, it may be possible to use supercoiling regulation for this, as it affects RNAP-promoter binding^61,62^. In that case, the location of the tandem promoters in the DNA (e.g. if in highly or weakly expressing topological domains^63,64^) could already suffice to influence the attenuation. Other influential variables are the durations of open complex formation and promoter escape, as they influence RNAP occupancy times of the promoter^65^. Changing both could even allow changing attenuation without changing transcription rates^11^.

Overall, our findings, including the model proposed to predict attenuation levels, should allow establishing a novel pipeline for engineering promoters (in tandem formation) whose overall kinetics can be fined-tuned within a relative wide dynamic state space. This pipeline should contribute to the development of novel components for complex synthetic genetic circuits with predictable behaviors, making their assembly faster and cheaper. Subsequently, the novel complex circuits could contribute to the engineering of bacterial strains whose metabolic tasks have minimal resource consumption, so as to improve the efficiency of bioindustrial processes.

## METHODS

### Bacterial strains, growth conditions, induction, and antibiotics

We used the *E. coli* strain DH5α-PRO (identical to DH5αZ1, here named “wild type”, WT)^13,14,66^. This strain produces the necessary regulatory proteins (LacI, AraC, and TetR) that tightly regulate each of our promoter constructs^13^.

First, chemically competent (CC) *E. coli* DH5α-PRO cells were prepared for plasmid transformations. For each strain, 10 ng of the plasmid DNA was mixed with 50 μL DH5α-PRO CC (1:10 ratio), and the mixture was incubated on ice for 30 min. Next, the mixture was kept at 42 °C in a water bath for 1 minute. Finally, 800 μL of LB medium was added to the mixture, which was kept at 37 °C under aeration at 250 RPM for 1 hour.

From each mixture, 200 μL was plated using the spread plate method on fresh LB agar plates (2%), prepared by supplementing with antibiotics (34 μg/mL chloramphenicol). Finally, the plates were kept overnight at 37 °C. The next day, three colonies were picked from each plate and inoculated in fresh LB medium, supplemented with antibiotics (34 μg/mL chloramphenicol). Afterwards, the cells were incubated at 30 °C overnight with shaking at 250 RPM. The resulting cells were diluted into fresh M9 media (0.03 O.D._600_) and were supplemented with 0.4% glycerol, amino acids, and vitamin solutions ^11,14^.

The control condition was M9 medium at 30°C. Cells were induced and given antibiotic treatment at the mid-exponential phase (O.D._600_ ≈ 0.25-0.3). Flow cytometry data was collected 180 minutes after induction. The spectrophotometry time series started immediately after induction and continued for 700 min.

The LacO_3_O_1_ promoter was induced with 1 mM IPTG, the TetA promoter was induced with 15 ng/mL aTc, and the BAD promoter was induced with 0.1% arabinose (induction curves in Figure S6)^14,66,67^. The promoters are referred to as P_LacO3O1,_ P_tetA,_ and P_BAD,_ respectively, for simplicity. In some experiments, cells were subjected to rifampicin (2.5 μg/mL, 1.25 μg/mL, 3.75 μg/mL and 5.0 μg/mL)^68^. Rifampicin (on its own) does not influence cell fluorescence (Figure S7).

### Spectrophotometry

We performed spectrophotometry to measure optical density at 600 nm (O.D._600_), which we used to monitor cell growth. We used a Biotek Synergy HTX Multi-Mode Reader. We also measured mean cell fluorescence. We used the excitation and emission wavelengths for mCherry (575/15 nm and 620/15 nm, respectively) with a gain of 50. Time series data was captured every 20 minutes. We normalized the fluorescence by the O.D._600_ to estimate the average single-cell fluorescence. For this, O.D._600_ curves were time shifted so that they reached 5% of their maximal O.D._600_ at the same time point., as in ^57^.

### Flow cytometry

We performed flow cytometry using an ACEA NovoCyte Flow Cytometer controlled by the software Novo Express V1.50. Cells were diluted 1:10000 into 1 ml of PBS, vortexed for 10 sec. For each condition, we performed 3 biological replicates, acquiring 50000 events for each replicate. We detected mCherry using the PE-Texas Red channel with excitation 561 nm and emission 615/20. We collected events at a flow rate of 14 μL/minute (core diameter of 7.7 μM). We have set a minimum detection threshold in FSC-H at 5000 to remove small particle interference. We also discarded the 1% of the highest PE-Texas Red-H intensities detected, to remove outliers.

### Fittings and statistical analysis

All best-fitting lines (e.g., Figure 2C) were obtained by linear regression (MATLAB). Fits of other functions (Figure 2D and Figure 8A) were obtained by the nonlinear least squares method (MATLAB). Their goodness of fit was estimated from R^2^ values.

Meanwhile, to determine if there is a significant correlation between any pair of variables, *y* and *x*, we used *t*-statistic with the null hypothesis that the slope of the best-fitting line (of a scatter plot between the variables) is not different from zero. For *p*-values smaller than 0.05, we reject the null hypothesis.

To identify outlier data points (in Figure 2C) we used an iterative procedure to discard outliers in linear correlations^40^. Data points (*x,y*) were classified as outliers when their vertical distance from the best linear fit is larger than the standard deviation of the distribution of *y* values (Figure S2). When applied, the process always converged in 1 iteration.

Finally, the associated standard error of the mean for calculations with fluorescence was calculated using an error propagation method. In general, if *z* = *f* (*x, y*), then the error of *z* is 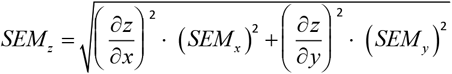, where 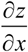 and 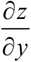 are the partial derivatives of *z* with respect to *x* and *y*, respectively.

### Stochastic simulations

We used stochastic simulations to simulate models used to estimate how changes in the dynamics of the promoters (e.g., RNAP occupancy time, repressor occupancy time, and frequency of dual RNAP fall-offs compared to one RNAP fall-off) influence the degree of attenuation in tandem promoter formations. The model is described in Supplementary Section S1. Simulations were performed using SGNS2^69^, a stochastic gene networks simulator whose dynamics follow the Stochastic Simulation Algorithm^70^. The time length of each simulation was set to 5×10^5^ seconds (with a sampling interval of 100 seconds). This time length sufficed to reach quasi-equilibrium in RNA numbers. The average behavior of each condition was characterized from 100 independent runs, as it always sufficed to obtain consistent results. Finally, for the various conditions, we assumed the parameter values in Supplementary Table S1 and the initial amounts of reactants in Supplementary Table S2.

## ACKNOWLEDGEMENTS

We thank Jane and Aatos Erkko Foundation [10-10524-38 to A.S.R.]; Sigrid Jusélius Foundation [230181 to A.S.R.]; Suomalainen Tiedeakatemia (to S.D. and to R.J.); EDUFI Fellowship [TM-21-11655 to R.J.]; Finnish Cultural Foundation [00222452 and 00232457 to I.S.C.B.]; Tampere University Graduate Program (to V.C.). The funders had no role in study design, data collection and analysis, decision to publish, or preparation of the manuscript. The authors also acknowledge the Tampere Flow Cytometry Facility for their service. Finally, Figures 1, 4, and 7A were created with BioRender.com.

## AUTHOR CONTRIBUTIONS

A.S.R. and V.C. conceived the study. A.S.R. supervised the study. V.C. designed the constructs and planned and executed most measurements. I.S.C.B. executed most data analysis. S.D. assisted in the design of the constructs and measurements. V.C. and R.J. contributed to the data analysis. A.S.R, R.J., and I.S.C.B. developed the model. I.S.C.B. implemented the model. A.S.R., V.C., and I.S.C.B. drafted all documents, which were revised by all co-authors.

