## Supplementary data for "Transcription attenuation in synthetic promoters in tandem formation"

**Supplementary Sections**

### S1. Stochastic model of tandem promoters

#### S1.1. Modeling single gene expression with two-step transcription initiation, repression, and induction mechanisms

We modelled active transcription initiation in *E. coli* as a two-step stochastic process [1]. Reaction S1 models an RNAP finding a promoter and committing to transcription initiation (completing the closed complex formation) at the stochastic rate constant $k_b$. Reaction S2 models the RNAP escape from the promoter (Pro), at the rate $k_{esc}$. This step accounts for open complex formation and promoter escape, which completes the initiation process [2,3]. This step also frees the promoter for new RNAPs.

After escaping, the RNAP quickly elongates ($RNAP_e$) and assembles the RNA coded by the gene. In the end, the RNA and RNAP are released (reaction S3). The number of nucleotides of the gene, *No. nuc.*, was set to 890 to match the average gene length of *E. coli* [4]. Instead, one could have set it to 708 bp, which is the length of the mCherry coding region (Results section "Assembly of the synthetic, non-overlapping tandem promoters in single-copy plasmids."), but this would not significantly affect the results.

Next, we model elongation as a single step, for simplicity (instead of multiple steps as in [5]). This does not influence mean elongation time lenght, only variance. We also included RNA degradation, in order to be able to estimate mean RNA numbers (reaction S4).

Finally, we added a repression mechanism. The repressors (Rep) binding rate to a free promoter is $k_r$ and the unbinding rate is $k_a$ (reactions S5). When Rep is bound, the promoter is unavailable to RNAP binding. The influence of repressors can be strongly reduced by inducers, *Ind*. These can directly bind/unbind repressors (at rates $k_{ind}$ / $k_{unind}$) (reaction S6). When bound by an inducer, the repressor cannot bind to the promoter.

$$Pro + RNAP \xrightarrow{k_t} Pro.RNAP \quad \text{(S1)}$$

$$Pro.RNAP \xrightarrow{k_{esc}} Pro + RNAP_e \quad \text{(S2)}$$

$$RNAP_e \xrightarrow{k_{elong}/No.nuc.} RNA + RNAP \quad \text{(S3)}$$

$$RNA \xrightarrow{k_d} \varnothing \quad \text{(S4)}$$

$$Pro + Rep \underset{k_a}{\overset{k_r}{\rightleftarrows}} Pro.Rep \quad \text{(S5)}$$

$$Rep + Ind \underset{k_{unind}}{\overset{k_{ind}}{\rightleftarrows}} Ind.Rep \quad \text{(S6)}$$

**S1.2. Modeling tandem promoters with transcriptional interference due to RNAP collisions leading to fall-offs with repression and induction mechanisms.**

To model tandem promoters, we adapted a model for operons with multiple promoters in tandem formation, recently published in [6]. First, we identify the promoters according to their position ($Pro^U$ and $Pro^D$, where U and D stand for upstream and downstream, respectively). Similarly, we identify elongating RNAPs ($RNAP_e$) and RNAs, depending on which promoter they start from. Given this, Reactions S7 and S8 model transcription initiation of the downstream and upstream promoter, respectively, as follows:

$$Pro^D + RNAP \xrightarrow{k_t^D} Pro^D.RNAP \quad \text{(S7)}$$

$$Pro^U + RNAP \xrightarrow{k_t^U} Pro^U.RNAP \quad \text{(S8)}$$

Next, reaction S9 allows the RNAP at $Pro^D$.RNAP to escape the promoter, making it available for new transcription events. Afterwards, the RNAP can complete elongation and the RNA be released (reaction S10):

$$Pro^D.RNAP \xrightarrow{k_{esc}^D} RNAP_e^D + Pro^D \quad \text{(S9)}$$

$$RNAP_e^D \xrightarrow{k_{elong}/No.nuc.} RNA^D + RNAP \quad \text{(S10)}$$

After forming $Pro^U$.RNAP complexes, the RNAP needs to escape the promoter (reaction S11):

$$Pro^U.RNAP \xrightarrow{k_{esc}^U} RNAP_e^U + Pro^U \quad \text{(S11)}$$

Then, the following events can occur to $RNAP_e^U$. First, if the downstream promoter is unoccupied, $RNAP_e^U$ can pass by and complete the RNA (reaction S12):

$$RNAP_e^U \xrightarrow{k_{elong}/No.nuc.} RNA^U + RNAP \quad \text{(S12)}$$

Contrarily, if the downstream promoter is occupied by an RNAP, $RNAP_e^U$ will collide with it. In that case: (i) either $RNAP_e^U$ will fall-off (reaction S13); or (ii) both RNAPs fall-off (reaction S14):

$$RNAP_e^U + Pro^D.RNAP \xrightarrow{k_c \cdot f_1} Pro^D.RNAP + RNAP \qquad \text{(S13)}$$

$$RNAP_e^U + Pro^D.RNAP \xrightarrow{k_c \cdot f_2} Pro^D + RNAP + RNAP \qquad \text{(S14)}$$

S13 and S14 are competing reactions, and one of them has to occur. We control their mean rate using a base rate constant ($k_c$). Meanwhile, we control their relative frequency by multiply $k_c$ with either $f_1$ or $f_2$, respectively, where: $f_1 + f_2 = 1$. Here, $f_1$ stands for the frequency with which only $RNAP_e^U$ falls-off, while $f_2$ stands for the frequency with which both RNAPs fall-off.

Noteworthy, at quasi-equilibrium, the RNAP collisions will affect mean RNA levels, only when causing RNAP fall-offs. Thus, since we are focusing on mean RNA levels, we do not model collisions that do not cause fall-offs. Also, we expect little differences in mean RNA levels at quasi-equilibrium between when the RNAP that falls-off is $RNAP_e^U$ or, instead, is the RNAP at the downstream promoter. Thus, we only model the first case (since the binding of $RNAP_e^U$ to the DNA should be comparatively weaker [7]).

Finally, we modeled external repression and activation of the promoters. Reaction S16 and S17 model the binding of specific repressors, $Rep^U$ and $Rep^D$, to the upstream and downstream promoters, respectively:

$$Pro^U + Rep^U \underset{k_a^U}{\overset{k_r^U}{\rightleftarrows}} Pro^U.Rep^U \qquad \text{(S16)}$$

$$Pro^D + Rep^D \underset{k_a^D}{\overset{k_r^D}{\rightleftarrows}} Pro^D.Rep^D \qquad \text{(S17)}$$

These repressors can be inactivated by the binding of their respective inducers, $Ind^U$ and $Ind^D$, respectively (reaction S18 for $Rep^U$ and reaction S19 for $Rep^D$), as follows:

$$Rep^U + Ind^U \underset{k_{unind}^U}{\overset{k_{ind}^U}{\rightleftarrows}} Ind^U.Rep^U \qquad \text{(S18)}$$

$$Rep^D + Ind^D \underset{k_{unind}^D}{\overset{k_{ind}^D}{\rightleftarrows}} Ind^D.Rep^D \qquad \text{(S19)}$$

Finally, we considered that, if the downstream promoter subject to repressors, the RNAP elongating from the upstream promoter could collide with a bound repressor. When occuring, similarly to when the downstream promoter is occupied by an RNAP, either $RNAP_e^U$ will fall-off (reaction S20, with frequency $f_r$) or the bound repressor will fall-off (reaction S21, with frequency (1- $f_r$)). For simplicitly, we do not model cases where both repressor and RNAP fall-off since, given the number of repressors and their binding and unbinding rate constants, this simplification did not influence the dynamics.

$$RNAP_e^U + Pro^D.Rep^D \xrightarrow{k_c \cdot f_r} Pro^D.Rep^D + RNAP \qquad \text{(S20)}$$

$$RNAP_e^U + Pro^D.Rep^D \xrightarrow{k_c \cdot (1-f_r)} Pro^D + Rep^D + RNAP_e^U \qquad \text{(S21)}$$

The RNAs produced in reactions S10 and S12 can decay via reactions S22 and S23, respectively:

$$RNA^U \xrightarrow{k_d} \varnothing \tag{S22}$$

$$RNA^D \xrightarrow{k_d} \varnothing \tag{S23}$$

Finally, all parameter values and supporting references are shown in Supplementary Table S1.

### Supplementary Figures

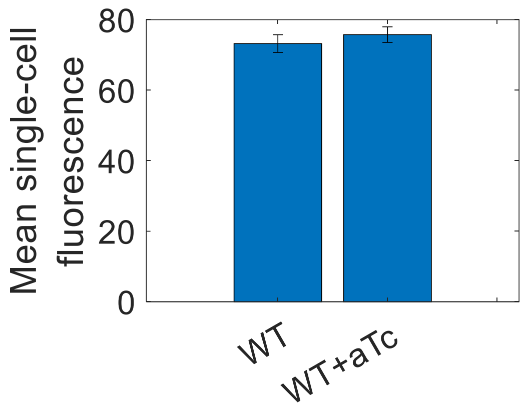


**Figure S1**: Single-cell fluorescence levels in arbitrary units. We used wild-type (WT) DH5α-Pro cells, absent of our plasmids, with and without the inducer, aTc. Fluorescence was measured 180 minutes (after adding aTc) at an O.D.$_{600}$ of 0.3. The error bars represent the SEM from 3 biological replicates.

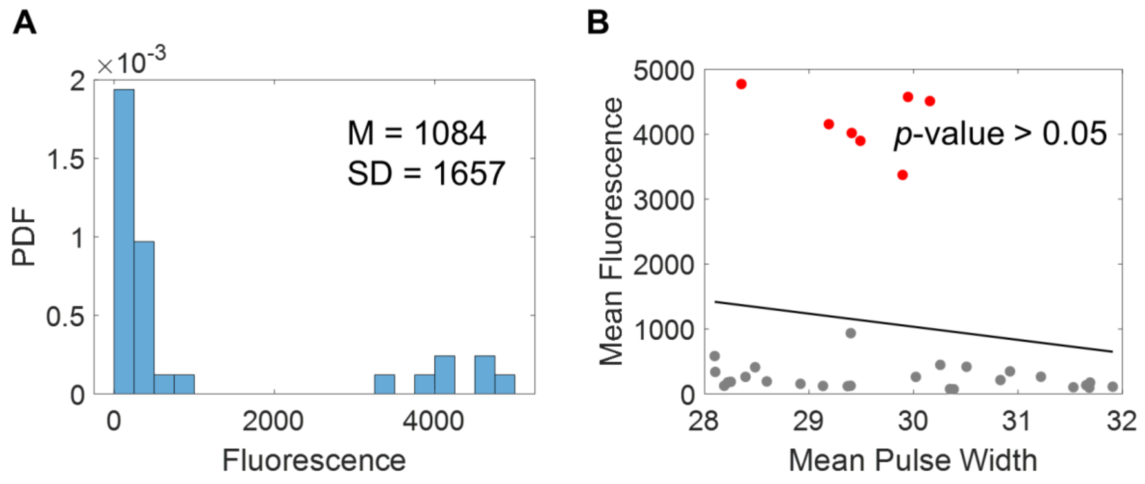


**Figure S2**: **(A)** Probability density function (PDF) of mean single-cell fluorescence levels. Also shown are the mean (M) and standard deviation (SD) of the distribution (top right). The SD is used as a threshold for the identification of outliers in Figure 2C in the main manuscript. **(B)** Scatter plot of the mean single-cell fluorescence plotted against the mean single-cell pulse width in each condition. The conditions are listed in Figure 2B in the main manuscript, including those classified as outliers (red balls). Also shown is a best fitting line. Visibly, the outliers (red points) affect the best-fitting line, causing it to not fit the bulk of the data points (grey). However, excluding them did not alter the conclusion, i.e., that one cannot conclude that the best fitting line fits better than a horizontal line ($p$-value > 0.05) (Methods section" Fitting and statistical analysis"). As such, we opted for not removing them from the line fitting.

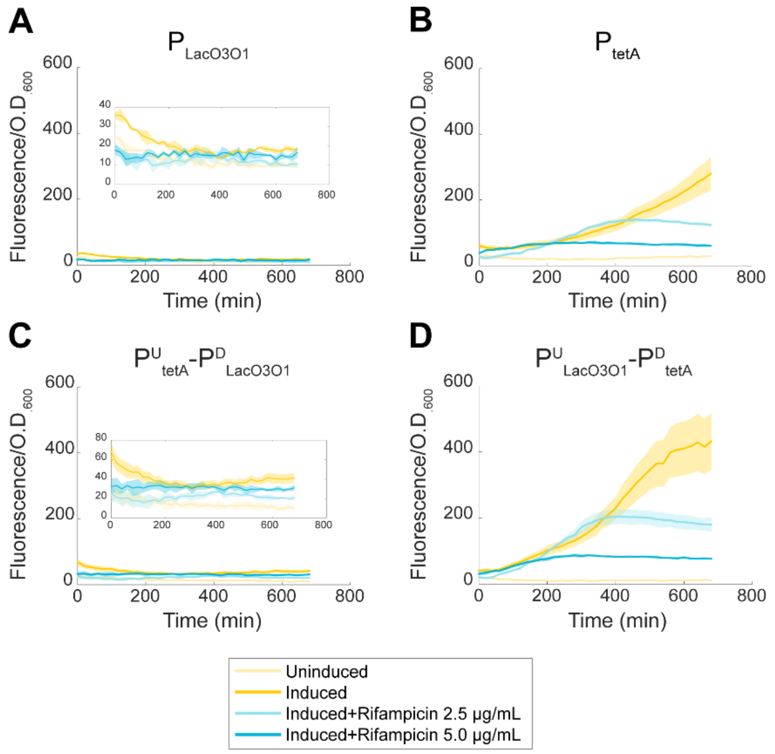


**Figure S3**: Average single-cell fluorescence levels of individual and tandem promoters. We measured population fluorescence levels (by spectrophotometry) and normalized them by the corresponding O.D.$_{600}$ to get a proxy for the average single-cell fluorescence levels (Methods section "Spectrophotometry"). Data for when uninduced, when fully induced (all promoters), and when fully induced while also subject to rifampicin (2.5 µg/mL and 5.0 µg/mL). The shaded areas represent the SEM from 3 biological replicates. "Induced" stands for when all promoters of the construct are fully induced. "Uninduced" stands for when no promoter is induced.

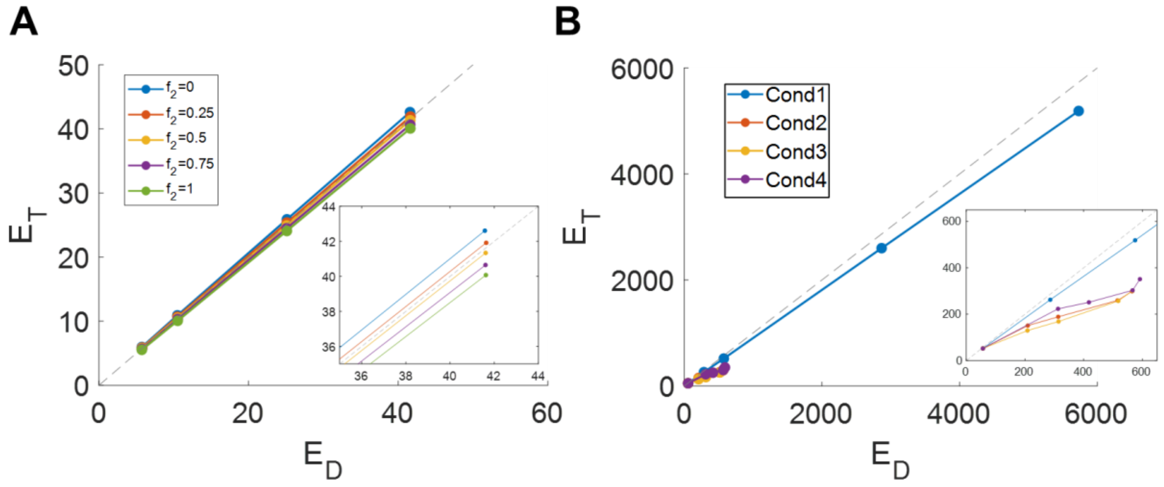


**Figure S4**: *In silico* expression levels of individual downstream promoters ($E_D$) and tandem promoters ($E_T$). **(A)** Effects of changing the RNAP escape rate from the downstream promoter ($k_{esc}^D$). Each colored line represents a different relative frequency ($f_2$) of both RNAPs falling-off upon colliding (as opposed to only one RNAP falling-off). **(B)** Effects of changing both the rates of RNAP binding and of RNAP escape from the upstream and from downstream promoter ($k_t^U$, $k_{esc}^U$, $k_t^D$, and $k_{esc}^D$). Each colored line represents a different set of parameter values modeling different conditions. Each condition ("Cond") is described in Supplementary Table S3. The dash line is the line of equality ($E_D$= $E_T$). Finally, the insets show zoomed areas of the plots, to better distinguish which data points are above and below the dashed line.

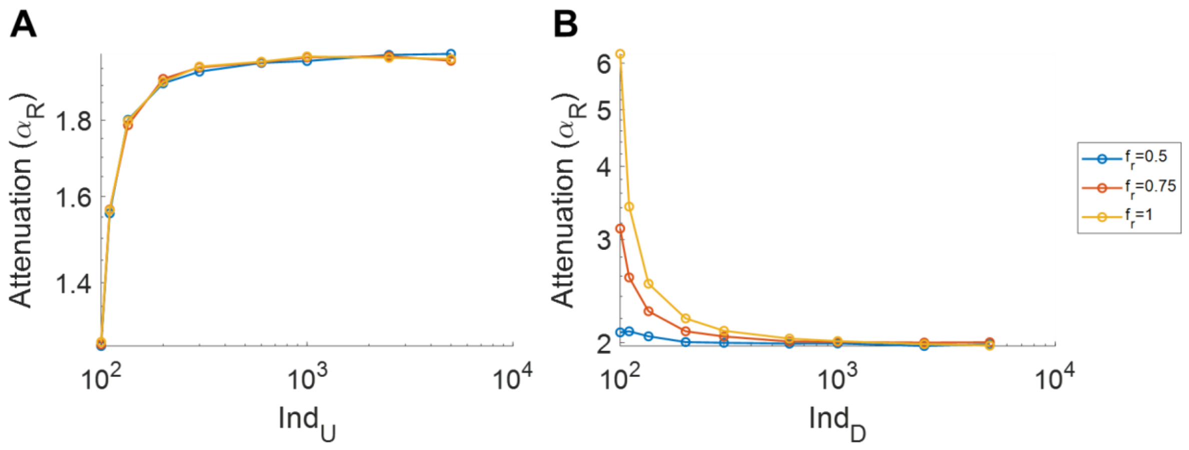


**Figure S5**: *In silico* estimation of the relative attenuation ($\alpha_R$) for tandem promoters when **(A)** the upstream promoter is gradually induced, and, when **(B)** the downstream promoter is gradually induced. The plots are in logarithmic scale. In (A) the downstream promoter is fully induced while in (B) the upstream promoter is fully induced. $f_r$ stands for the relative frequency with which an RNAP from the upstream promoter falls-off upon colliding with a repressor bound to the downstream promoter (as opposed to the repressor falling-off). For $f_r$ = 1, it is always the RNAP that falls-off with the collision.

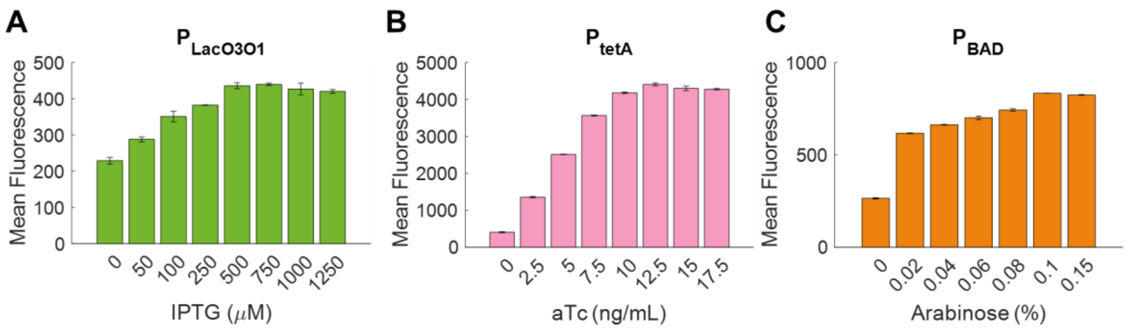


**Figure S6:** Induction curves of $P_{LacO3O1}$ (inducible by IPTG), $P_{tetA}$ (inducible by aTc), and $P_{BAD}$ (inducible by arabinose) in individual formation, as measured by single-cell fluorescence. The mean and standard error of the single-cell fluorescence were extracted from 3 biological replicates in each condition. In all cases, the strongest inducer concentration resulted in a minor decrease in fluorescence. For this reason, the previous condition was used as being "maximum induction".

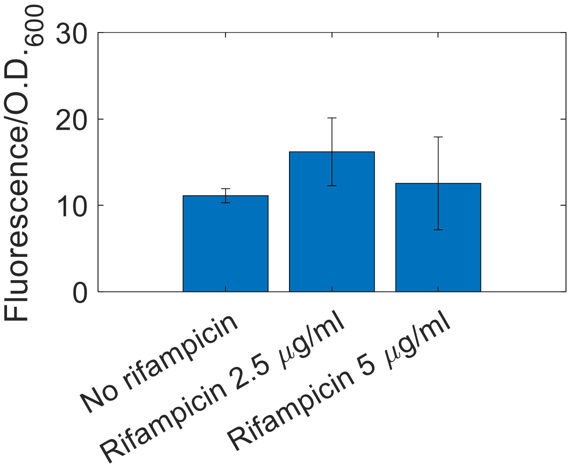


**Figure S7**: Fluorescence levels of DH5α-Pro cells (WT) absent of our plasmids when subject to rifampicin. Population fluorescence levels (spectrophotometry) are normalized by the corresponding O.D.$_{600}$, to be used as a proxy for average single-cell fluorescence (Methods section "Spectrophotometry"). We measured 180 minutes after adding rifampicin, when reaching O.D.$_{600}$ of 0.3. The error bars represent the SEM from 3 biological replicates.

### Supplementary Tables

**Table S1:** Parameter values of the model.

| Parameter | Value | Reference |
|---|---|---|
| $k_t^D$ and $k_t^U$ | 0.0013 $s^{-1}$ | [1] |
| $k_{esc}^D$ | 0.0052; 0.0104; 0.0347; 0.104 $s^{-1}$ | Scaled to cover most of the possible state space. These values were used to test corresponding occupancy times of 87%, 75%, 50%, and 30% of the simulation time, approximately. |
| $k_{esc}^U$ | 0.0052 $s^{-1}$ | [1] |
| $k_{elong}$ | 42 nucleotides/s | [8] |
| *No. nucleotides from the TSS to the transcription termination site.* | $Pro^D$: 890 nucleotides $Pro^U$:1030 nucleotides | Average values estimated from [4,9], using the classification of tandem promoters from [9]. |
| $k_c$ | 1/3 $s^{-1}$ | Base value of the expected time that it will take an RNAP elongating from the upstream promoter to find (if present) an RNAP bound to the downstream promoter. Estimated using $k_{elong}$ and the average distance of 140 nucleotides between natural TSSs [9]. In our constructs, this distance is 150 nucleotides. |
| $f_2$ | 0; 0.25; 0.5; 0.75; 1 | |
| $k_r^D$ and $k_r^U$ | 1 $s^{-1}$ | [10] |
| $k_a^D$ and $k_a^U$ | 0.1 $s^{-1}$ | [10] |
| $k_{ind}^D$ and $k_{ind}^U$ | 1 $s^{-1}$ | For simplicity, Rep molecules are equally likely to bind to an inducer as to the promoter. |
| $k_{unind}^D$ and $k_{unind}^U$ | 0.1 $s^{-1}$ | |
| $f_r$ | 0.5; 0.75; 1 | |
| $k_d^U$ and $k_d^D$ | 0.00082 $s^{-1}$ | [11] |

**Table S2:** Initial amounts of reactants in simulations. The initial values of all reactants not shown are set to 0.

| Reactant | Initial value | Reference |
|---|---|---|
| RNAP | 40 | [1] |
| $Rep^U$ and $Rep^D$ | 100 | [10] |
| $Ind^U$ and $Ind^D$ | 0; 100; 110; 135; 200; 600; 1000; 2500; 5000 | Set to cover most of the possible state space of repression strength. |

**Table S3:** Parameter values of the different conditions in the plots in Figure S4B.

| Condition | Description | **($k_t^U$ ; $k_t^D$) ($s^{-1}$)** | **($k_{esc}^U$ ; $k_{esc}^D$) ($s^{-1}$)** | **$f_2$** |
|---|---|---|---|---|
| 1 | Increase all rates by the same amount | (0.013; 0.013)<br>(0.065; 0.065)<br>(0.13; 0.13)<br>(0.65; 0.65)<br>(1.3;1.3) | (0.052; 0.052)<br>(0.26; 0.26)<br>(0.52; 0.52)<br>(2.6; 2.6)<br>(5.2; 5.2) | 1 |
| 2 | Increase $k_{esc}^U$ and $k_{esc}^D$ by the same amount | (0.013; 0.013) | (0.052; 0.052)<br>(0.26; 0.26)<br>(0.52; 0.52)<br>(2.6; 2.6)<br>(5.2; 5.2) | 1 |
| 3 | Increasing both $k_{esc}^U$ (strong increase) and $k_{esc}^D$ (weak increase) | (0.013; 0.013) | (0.052; 0.052)<br>(0.52; 0.26)<br>(1.04; 0.52)<br>(5.2; 2.6)<br>(10.4; 5.2) | 1 |
| 4 | Increasing both $k_{esc}^U$ (weak increase) and $k_{esc}^D$ (strong increase) | (0.013; 0.013) | (0.052; 0.052)<br>(0.26; 0.52)<br>(0.52; 1.04)<br>(2.6; 5.2)<br>(5.2; 10.4) | 1 |

**Table S4:** Values of the coefficients of the best fittings in the main manuscript. We only present linear fits for correlations considered to be significant (Methods section "Fittings and statistical analysis").

| Figure | Fitting formula | Coefficient values |
|---|---|---|
| Figure 2D | $y = \frac{C}{x} + n$ | $C$ = 231<br>$n$ = 1.02 |
| Figure 5A$_1$ ($P_{tetA}$) | $y = \frac{a}{1+e^{-b\cdot(x-c)}}$ | $a$ = 4370<br>$b$ = 0.49<br>$c$ = 4.31 |
| Figure 5A$_1$ ($P^U_{tetA}$- $P^D_{LacO3O1}$) | $y = \frac{a}{1+e^{-b\cdot(x-c)}}$ | $a$ = 404.5<br>$b$ = 0.19<br>$c$ = -2.37 |

| | | |
|---|---|---|
| Figure $5A_2$ | $y=\frac{a\cdot x+b}{x+c}$ | $a$ = 15.65<br>$b$ = 14.52<br>$c$ = 4.83 |
| Figure $5B_1$ ($P_{LacO3O1}$) | $y=\frac{a}{1+e^{-b\cdot(x-c)}}$ | $a$ = 428<br>$b$ = 0.01<br>$c$ = -17.7 |
| Figure $5B_1$ ($P^{U}_{tetA}$- $P^{D}_{LacO3O1}$) | $y=\frac{a}{1+e^{-b\cdot(x-c)}}$ | $a$ = 388.7<br>$b$ = 0.009<br>$c$ = 47.89 |
| Figure $5B_2$ | $y=\frac{x+a}{b\cdot x+c}$ | $a$ = 283.4<br>$b$ = 0.10<br>$c$ = 9.52 |
| Figure $7C_1$ | $y=a\cdot x+b$ | $a$ = 0.04<br>$b$ = 1.05 |
| Figure $7C_2$ | $y=a\cdot x+b$ | $a$ = 4.68<br>$b$ = 10.72 |
| Figure 8A ($f_2$ = 0) | $y=\frac{a\cdot x+b}{x+c}$ | $a$ = 1.00<br>$b$ = -0.005<br>$c$ = 20.16 |
| Figure 8A ($f_2$ = 0.25) | $y=\frac{a\cdot x+b}{x+c}$ | $a$ = 0.98<br>$b$ = -0.02<br>$c$ = 19.81 |
| Figure 8A ($f_2$ = 0.5) | $y=\frac{a\cdot x+b}{x+c}$ | $a$ = 0.95<br>$b$ = 0.11<br>$c$ = 20 |
| Figure 8A ($f_2$ = 0.75) | $y=\frac{a\cdot x+b}{x+c}$ | $a$ = 0.93<br>$b$ = 0.01<br>$c$ = 19.51 |
| Figure 8A ($f_2$ = 1) | $y=\frac{a\cdot x+b}{x+c}$ | $a$ = 0.91<br>$b$ = 0.05<br>$c$ = 19.26 |
| Figure 8B ($f_2$ = 0) | $y=a\cdot x+b$ | $a$ = 0.004<br>$b$ = 1.06 |
| Figure 8B ($f_2$ = 0.25) | $y=a\cdot x+b$ | $a$ = 0.005<br>$b$ = 1.09 |
| Figure 8B ($f_2$ = 0.5) | $y=a\cdot x+b$ | $a$ = 0.005<br>$b$ = 1.10 |
| Figure 8B ($f_2$ = 0.75) | $y=a\cdot x+b$ | $a$ = 0.005<br>$b$ = 1.12 |
| Figure 8B ($f_2$ = 1) | $y=a\cdot x+b$ | $a$ = 0.005<br>$b$ = 1.14 |
